# Method and timing of rhizobacteria inoculation to plant roots alters success and strength of aphid suppression

**DOI:** 10.1101/2024.07.30.605806

**Authors:** Sophie Blenkinsopp, Milo Henderson, Sharon E Zytynska

**Affiliations:** Department of Evolution, Ecology, and Behaviour. Institute of Infection, Veterinary and Ecological Sciences. University of Liverpool, Crown Street, Liverpool, L69 7ZB, UK; Plant and Soil Science, Scotland’s Rural College, Peter Wilson Building, King’s Buildings, W Mains Rd, Edinburgh EH9 3JG, UK

## Abstract

Insect pests cause substantial yield losses globally, necessitating novel pest control approaches beyond chemical pesticides. Rhizobacteria, beneficial root-associated bacteria, present a promising alternative by enhancing plant growth and defences against pests. This study explores the use of sodium alginate encapsulation for precise inoculation of two rhizobacteria, *Acidovorax radicis* and *Bacillus subtilis*, to suppress aphid populations on barley (*Hordeum vulgare*). We optimized a method using 4mm diameter wet-matrix alginate beads for controlled bacterial delivery directly to plant roots. Two experiments evaluated the impact of inoculation methods and timing on aphid suppression. Results indicated that rhizobacteria inoculation via alginate beads at root emergence significantly reduced aphid reproductive output, outperforming seed soaking methods. This suggests that more consistent bacterial establishment and prolonged release from alginate beads enhances plant defence priming. Additionally, alginate bead inoculation demonstrated effective long-term bacterial viability following storage at 4° C for eight months, supporting potential field application. Our findings highlight the potential of alginate bead-encapsulated rhizobacteria for aphid suppression on barley, but emphasizing the need for precise inoculation timing and placement. This approach offers a robust method for empirical research and practical agricultural application, paving the way for sustainable pest management strategies. Future work should focus on optimizing formulations and understanding plant-microbe interactions to enhance field efficacy.

## Introduction

Insect pests cause significant yield losses worldwide and there is an immediate need to develop novel approaches that complement or replace chemical pesticides (Douglas 2018). Emerging pesticide resistance of many arthropod species (Whalon, Mota-Sanchez & Hollingworth 2008) strengthens the need and urgency for establishing and optimising novel methods for controlling insect populations. A promising approach is to apply plant-beneficial root-associated or rhizobacteria strains to crop roots, which can promote plant growth and induce plant defences against insect pests resulting in increased yields (Berendsen, Pieterse & Bakker 2012; Pieterse *et al*. 2014; Trivedi *et al*. 2021). A fast-growing body of research aims to identify how individual strains provide benefits (Blake, Christensen & Kovács 2021; Pieterse *et al*. 2021), but also to develop synthetic communities (SynComs) containing multiple rhizobacteria strains to enhance establishment and functional stability in field soils (Song *et al*. 2020; Shayanthan, Ordoñez & Oresnik 2022).

Microbial inoculation of plants aims to prime a plant’s defences before insect attack, enabling a stronger, faster response when they are eventually attacked by a pest (Conrath 2009). Several strains of rhizobacteria, and multi-strain rhizobacteria communities, have shown strong effects on reducing both chewing and sucking herbivore populations (Zytynska, Parker & Sanchez-Mahecha 2024). Rhizobacteria inoculation methods vary from seed soaking, seedling or root soaking, soil inoculation, and leaf spraying. Bashan (1986) described a method for using sodium alginate encapsulation of *Azospirillum brasilense* and *Pseudomonas* sp. strain 84313 for slow-release of bacteria that promote growth of wheat plants. Since then, several studies have continued to explored the use of this method to encapsulate plant-growth-promoting rhizobacteria (Hernández-Montiel *et al*. 2017), sustain bacteria viability under plant drought stress (Souza-Alonso *et al*. 2021), to encapsulate semio-chemicals for attracting natural enemies to crop plants (Heuskin *et al*. 2012), and in human probiotics to improve viability during gastrointestinal digestion (Cedran, Rodrigues & Bicas 2021). Alginate encapsulation of bacteria, using very small beads, has also successfully been used for seed coating of maize in combination with gum arabic (Berninger, Mitter & Preininger 2016). Effective inoculation of a plant with beneficial rhizobacteria needs to guarantee colonisation, establishment and persistence by the bacteria in the rhizoplane next to the root surface. Once there is inoculum in the rhizosphere, the secretion of root exudates will enable the bacteria to colonise the roots (Hu *et al*. 2018). Therefore, inoculation methods must ensure there are viable bacteria present ready to make associations in the soil when roots emerge from seeds for effective inoculation. Sodium alginate encapsulation of rhizobacteria is a promising method for enabling precise inoculation of bacteria to experimental plants, supporting continued research, and for large-scale production efforts to support field trials and eventually wide-spread agricultural use (Bashan *et al*. 2002). Co-entrapment of a rhizobacteria and mycorrhizal fungi led to increased root colonisation in potato plants, indicating the broad applicability of this method (Lojan *et al*. 2017). Finally, Trivedi and Pandey (2008) demonstrated successful recovery of *Bacillus subtilis* and *Pseudomonas corrugata*, immobilized in a sodium alginate based formulation, after 3 years of storage in 4 °C in the order of 10^8^ cfu g^−1^.

In this study, we explored the use of bacterial encapsulation of two rhizobacteria, *Acidovorax radicis* and *Bacillus subtilis*, that suppress aphid population growth on barley (*Hordeum vulgare*) (Álvarez-Lagazzi *et al*. 2021; Xi, Dean & Zytynska 2024). The size and formulation of beads can impact production, stability, and bacteria viability (John *et al*. 2011). Here, we optimised a method using large (4mm diameter) wet-matrix beads for individual plant inoculation with a controlled amount of bacteria, directly to plant roots, and facilitating ongoing research projects. We tested a range of additional components to the sodium alginate beads, then performed two experimental studies to assess the impact of inoculation method and timing of inoculation on the success of microbe-induced aphid suppression in the plants.

## Materials and Methods Study system

We individually inoculated two bacteria (*Acidovorax radicis* N35, *Bacillus subtilis* B171, or a control solution) onto roots of barley plant (*Hordeum vulgare*, varieties *Barbarella* and *Firefoxx* Elsoms Seeds) and challenged the plants with aboveground sap-feeding aphids (*Sitobion avenae*, genotype STC22p collected from a field trial at Stockbridge Technology Centre, Selby in 2022 and maintained as a clonal line in the lab on *Curtis* barley). All plants were germinated and grown in Levington’s Advance F1 low nutrient compost.

*Acidovorax radicis* bacteria were grown on nutrient agar plates [Difco™ General purpose Nutrient Broth 8g/litre plus 15g agar] at 30°C for 5 days, and harvested using cell scrapers into 10mM MgCl_2_. *Bacillus subtilis* was grown in 5ml cultures of nutrient broth, with continuous shaking at 30°C for 3 days, then transferred to 500ml flasks, containing 100ml of nutrient broth, and shaken at 30°C overnight. The bacteria were harvested via centrifugation and cleaned using 10mM NaCl. Both bacteria were resuspended in 10mM MgCl_2_ at OD_600_=2.0 (approx. 108 colony-forming units (cfu/ml)). As a no-bacteria control we used 10mM MgCl_2_.

### Seed and alginate bead preparation

All experimental seeds were surface sterilised with 4% bleach solution and washed thoroughly with tap water. Seeds were soaked in either bacterial inoculant or control solution (MgCl_2_) for 2 hours before transfer to potting soil for germination.

Alginate beads were made by mixing 5% sodium alginate powder into the bacterial solution (final equivalent concentration of OD_600_=2.0), and then dropped into 5% CaCl solution using a 10ml syringe from a height of 5cm. Droplets form beads in the CaCl solution, and consistent pressure on the syringe ensured individual beads of comparable size were made. The beads were stored overnight in CaCl to ensure matrix formation before being stored in minimal amount of CaCl to avoid drying until use. We tested several additions to the beads (to potentially enhance structure and bacteria viability) including glycerol (5%), nutrient broth (75%), nutrient broth with agar, spirulina (5%), and biochar (5%). For each addition, the final concentration of bacteria was constant at the equivalent of OD_600_=2.0.

For the final method (nutrient broth addition), we optimised the protocol to first add sodium alginate to autoclaved nutrient broth to facilitate mixing; a short burst of maximum 1 minute heating in the microwave was also beneficial to avoid lumps, as well as leaving overnight (or longer) in a cool room. To maximise bacterial concentration and growth, experimental beads were placed into a flask of nutrient broth and shaken overnight at 30°C for secondary bacterial multiplication, then removed and washed in sterile water. To estimate bacterial concentrations, alginate beads were dissolved in 0.2M potassium phosphate buffer pH 6.8 and released bacteria counted using dilution series, testing original (no secondary multiplication) beads as well as those that did. A set of beads were stored at 4C for long-term viability measures (up to 8 months).

### Experimental design and setup

All experiments are fully-factorial and were set up using a randomised complete block design to reduce variation across replicates.

Our first experiment was conducted in a glasshouse in April 2022 (average temperature of 20°C, with a minimum of 18°C and occasional peaks of higher temperatures depending on the outside weather), with six replicates per treatment. Here we compared seed soaking and bead inoculation (after seed germination), and a combined treatment of seed soaking plus bead inoculation, on aphid numbers. Seeds (*Barbarella* variety) were soaked in bacterial (*A. radicis* or *B. subtilis*) or control solution (as above) and germinated in soil using seed trays. Seedlings were transferred into individual 9 cm diameter pots on day 6, measuring plant shoot and root length, and beads were added by placing the bead under the soil surface next to the plant roots. On day 9, two 3rd-instar aphids were transferred to the plant using a fine paintbrush, and pots were covered by mesh cages to avoid aphid movement among plants. The total number of aphids (separated by reproducing adults and nymphs) were counted after two weeks, and plant shoot and root length were measured.

The second experiment was designed to test the effect of timing of bead inoculation, using barley variety *Firefoxx*. This experiment was run in a control temperature growth cabinet (20°C, 65% RH 16:8 light:dark), with eight replicates per treatment. We had five inoculation treatments for the two bacteria (*A. radicis* or *B. subtilis*) and the control treatment: (i) seed soaking, day 0, (ii) bead inoculation at seed sowing, day 0, (iii) bead inoculation after germination, day 5, (iv) bead inoculation at aphid infestation, day 7, and (v) after aphid infestation, day 11. Seeds were prepared as above and germinated in individual 9 cm diameter pots, shoot height was measured throughout the experiment. Beads were added to the seed/roots just under the soil surface. For this experiment we transferred two adult aphids to the plant (day 7), allowed them to reproduce for 3 days and then removed all aphids except two 1st instar offspring. These were allowed to develop to adults, and then offspring were individually removed (leaving the adults) every 2-3 days to allow a complete count of fecundity and adult survival (only two aphids survived beyond 30 days).

### Data analysis

All data were analysed using R v4.3.0 in R Studio v2023.03.0. For the first experiment, we used linear models to estimate the effect of bacterial treatment (*A. radicis, B. subtilis*, control) and inoculation method (seed, bead, or double inoculation) on the number of offspring and total number of aphids after two weeks. Replicate was included as a blocking factor, and plant growth as covariates. We also included explanatory factors to explore effects of single vs double inoculation (averaging across method of inoculation) and effects of bead vs seed (averaging across number of inoculations, and allowing an interaction between these to be tested). Due to the strong differences between the two bacteria, we also analysed these separately to explore bacteria-specific effects. While all statistical analyses were on aphid number data, we present these values relative to the control for the figures to highlight the aphid suppression effects where points are below the zero line.

For the second experiment, we explored the data in three main ways: (i) totals across the experiment, (ii) offspring production over time (cumulative and daily reproduction per surviving adult), and (iii) survival of adult aphids. For the totals, we calculated the total number of offspring produced by the adult aphids and used linear models (normal error distribution as this fitted model assumptions) to analyse the effect of bacterial treatment and inoculation method/timing on total fecundity. To simplify the interpretation, we calculated the number of aphids on bacterial-inoculated plants compared to their relevant controls and used linear models to estimate the difference in suppression effect (if any) across the inoculation methods. We also calculated the number of adult reproductive days and lifespan, but found no significant effects. For the data over time, we used linear mixed effects (repeated measures) models with pot as a random factor and including a quadratic term to account for non-linearity, to measure the effect of bacterial treatment and inoculation method/timing on the number of offspring produced. For one model we used the cumulative number of offspring produced, highlighting the population level impact on growth rates, and another we adjusted for the number of offspring per surviving adult per day (since offspring number was counted every 2-3 days). Lastly, aphid survival was analysed using the Cox proportional hazards regression model, using the R library (survival), as a response to bacterial treatment and inoculation method.

## Results

### Bead composition and storage

We tested a set of components aimed to stabilise the beads and promote bacterial persistence, based on previous literature. We found that (i) glycerol was unsuitable since beads did not form well and (ii) agar addition helped bead structural integrity but increased bead size variation and were more prone to containing air pockets making them less suitable for experimental work; (iii) spirulina, to support bacterial growth, led to increased contamination; (iv) biochar, to support bacterial growth, was too coarse for the small beads; and, (v) nutrient broth addition, to support bacterial growth, was easy to handle and allowed for good bead formation. In wet soil, bead breakdown occurred over the course of 24 hrs, with no remaining bead structure after 48 hrs.

Following, we continued to use the simple method of adding nutrient broth (NB) to the sodium alginate before mixing with bacterial solution. With an average bead volume of 0.1ml (1 ml of solution made approx. 10 beads), each bead was expected to initially contain 2.5 × 10^7 CFU/ml. Using dilution series, we found individual beads to contain higher concentration of bacteria than estimated (*A. radicis*: (2.7 +/-1.3) x 10^10 CFU/ml and *B. subtilis*: (6.8 +/-1.6) x10^9 CFU/ml). The bacteria remained viable for at least eight months of storage at 4°C with higher yield recovered than from fresh beads (*A. radicis*: (2.6 +/-1.2) x 10^14 CFU/ml and *B. subtilis*: (4.2 +/-1.9) x10^14 CFU/ml). Following an overnight multiplication period (at 30°C), we were able to recover an even higher bead bacterial concentration (*A. radicis*: (2.9 +/-1.3) x 10^15 CFU/ml and *B. subtilis*: (2.2 +/-1.1) x10^15 CFU/ml).

### Effect of bacterial inoculation method and timing on aphid suppression

In a first experiment, we observed variation in the strength of aphid suppression by the two bacteria dependent on inoculation method, with higher aphid suppression observed on *A. radicis* plants when inoculated via a bead than via seed soaking (F_2,29_=3.34, P=0.049; Fig.1). *Bacillus subtilis* plants exhibited stronger aphid suppression after seed soaking than bead inoculation (F_1,16_=7.48, P=0.014; Fig.1), partially driven by reduced variability among replicates. We also found that there was a loss of aphid suppression in the double inoculated *B. subtilis* plants compared to either seed-soaked or bead-inoculated plants (single vs double: F_1,31_=4.06, P=0.052; Fig.1). Lastly, we observed high levels of variation among replicates, which was not explained by pot location in the experimental space (F_1,26_=2.56, P=0.122) and only minimally influenced by variation in plant shoot growth (F_1,26_=4.07, P=0.053; used as a covariate in models).

**Figure 1.**
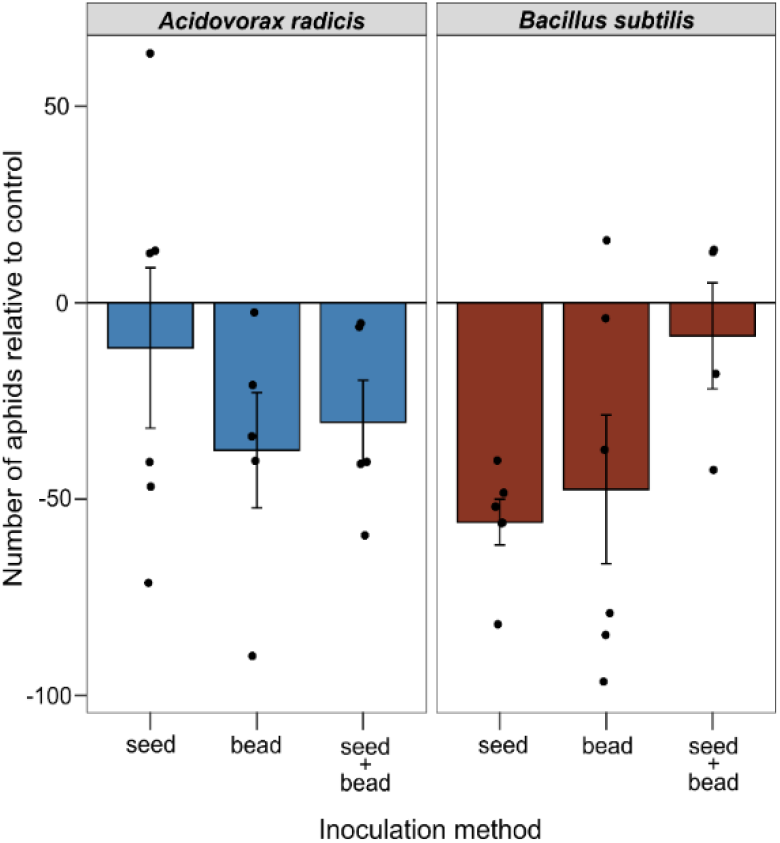
Number of aphids on inoculated plants relative to controls. We inoculated plants with rhizobacteria (A. radicis or B. subtilis) using seed soaking, bead addition, or both as methods of inoculation. Error bars ±1SE.

Our second experiment examined the effect of seed soaking compared to bead inoculation at different time points in the experiment (Fig. 2). For the control plants, there was no difference in total aphid number across inoculation method (F_4,31_=0.34, P=0.846), but we observed significant variation in aphid number across the inoculation methods for bacteria-inoculated plants (F_4,68_=2.88, P=0.029). By comparing relative effects compared to control plants, we observed a strong suppression effect of bacteria on aphids when the plants had been inoculated just after germination, ‘bead at germination’ (full model F_4,61_=6.20, P<0.001, treatment posthoc: t=3.70, P<0.001; Fig. 2a). There was no significant difference in total aphid numbers between the two rhizobacteria species, indicating a general rhizobacteria effect (F_1,61_=0.31, P=0.580). In addition, aphids were reduced on plants inoculated with B. subtilis after initial aphid infestation (‘bead after aphids’ treatment posthoc: t=2.33, P=0.025); Fig. 2a).Similar to the previous experiment, we observed high levels of variation within the treatments that could not be attributed to location in the growth chamber (rep: F_7,61_=0.75, P=0.628) or plant growth (F_1,60_=1.21, P=0.294).

**Figure 2.**
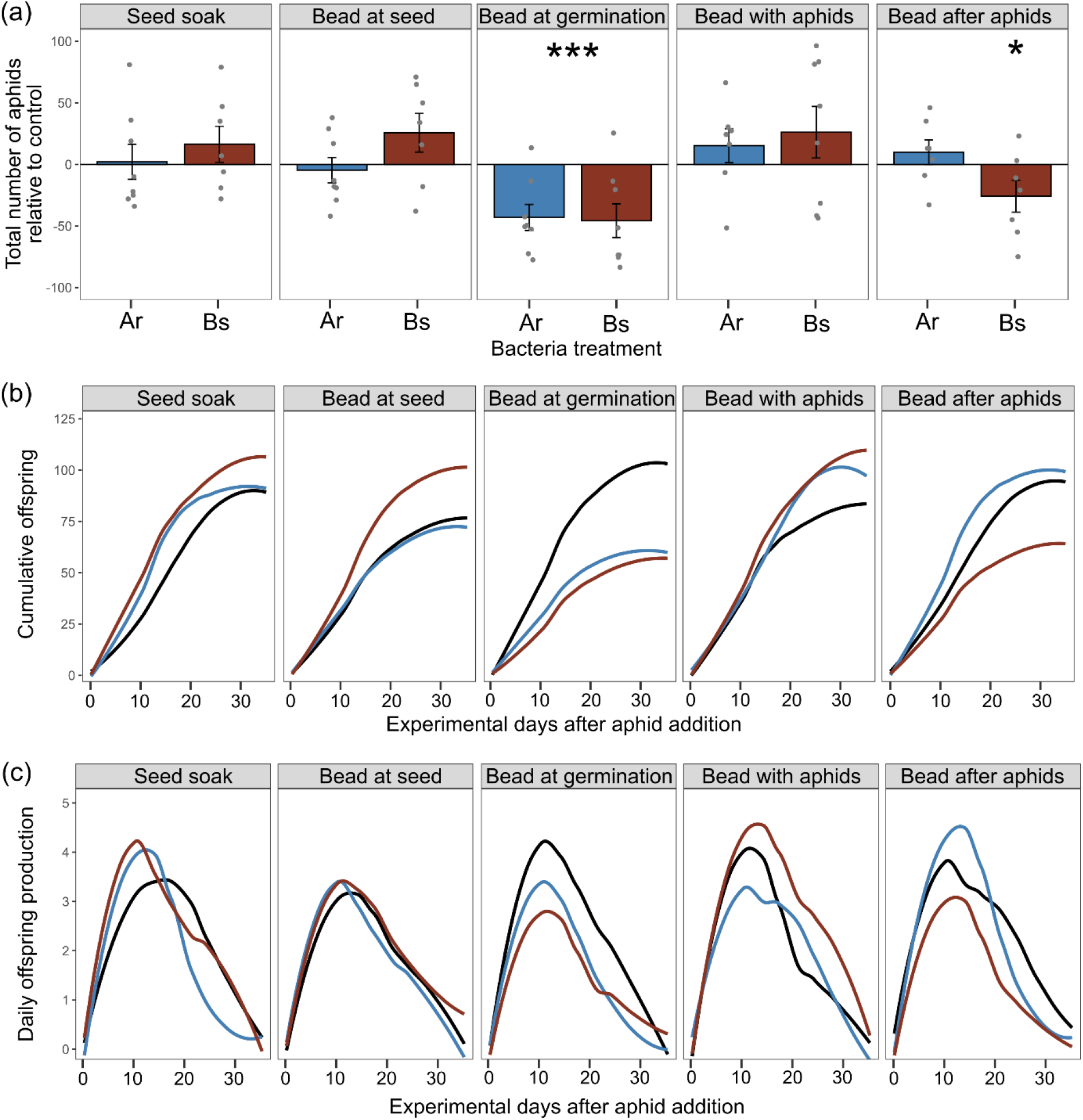
Aphid population growth over different inoculation treatments. We inoculated plants with rhizobacteria (*A. radicis* or *B. subtilis*) using seed soaking, bead addition (at seed sowing, at germination when roots just emerged, when we added aphids, or 4 days after initial aphid addition). (a) Total number of aphids on inoculated plants relative to control plants; Error bars ±1SE, (b) cumulative number of offspring and (c) daily offspring production by surviving adult aphids over the experimental period, smooth line fitted from loess model fit across 8 replicates; lines colour: control (black), *A. radicis* (blue), *B. subtilis* (red).

Aphid cumulative fecundity, as analysed over time, was significantly affected by bacterial treatment dependent on inoculation method (bacteria x inoculation, X^2^=16.84, df=8, P=0.031; Fig 2b). Aphid daily reproduction was significantly reduced by *B. subtilis* inoculation (X^2^=4.01, df=1, P=0.045; Fig 2c) with significant variation across inoculation methods (X^2^=10.59, df=4, P=0.032), and as an interaction between bacteria treatment and inoculation method (X^2^=9.09, df=8, P=0.059). We did not observe the same pattern for *A. radicis* inoculated plants, where the strongest effect on daily reproduction was for the subset of plants inoculated with a ‘bead at germination’ (X^2^=3.04, df=1, P=0.081; Fig 2c). Bacterial inoculation did not reduce aphid survival (X^2^=0.783, df=1, P=0.376) and we observed no difference in survival across the method of inoculation (X^2^=0.155, df=4, P=0.818), even for the subset of data from ‘bead after germination’ for both bacteria (X^2^=0.839, df=2, P=0.658).

## Discussion

Our results show that rhizobacteria inoculation of barley plants can suppress aboveground aphids, primarily through reduction in reproductive output. Yet the success and strength of effect was dependent on the method and timing of inoculation. Our data showed more consistent success when using alginate-bead encapsulation of bacteria inoculated to plants upon root emergence. The first experiment showed significant suppression of aphids by *Bacillus subtilis* following both seed soaking and bead addition but only bead inoculation was successful in the second experiment. Previous work (Xi, Dean & Zytynska 2024) showed significant effects of *A. radicis* on aphid suppression using seed soaking, but we did not observe the same pattern here, with only significant reduction from bead inoculation. The large variation in responses within treatments may explain this: successful establishment of bacteria leads to plant suppression of aphids, but this does not occur in all replicates. Seed soaking requires the bacteria to persist until root emergence (∼3 days for barley), and thus may lead to greater variation in successful inoculations. With bead inoculation at root emergence, we observed more consistent (and thus statistically significant) reduction in aphid reproduction and population growth across replicates. Better establishment of the bacteria in these replicates due to the presence of roots at the time of inoculation, and the slow release of bacteria over bead breakdown period of up to 48 hours may contribute to this (Bashan 1986; Hernández-Montiel *et al*. 2017). Bead inoculation can therefore support continued delivery of bacteria until root-interactions are established. For both empirical experimental research and field delivery, this is of additional benefit under adverse climate conditions, e.g. alginate encapsulation supported bacteria viability under drought conditions (Souza-Alonso *et al*. 2021).

We found suppression effect on aphid population growth occurred via reduction in reproductive output rather than survival of aphid individuals. This is in agreement with data from our recent meta-analysis that showed stronger rhizobacteria-mediated effect on sucking-insect fecundity than survival (Zytynska, Parker & Sanchez-Mahecha 2024). Our experimental design inoculated plants either before aphid infestation (at least 2 days before), at the same time, or four days after aphid infestation. The lack of effect when plants are inoculated at the same time as aphid infestation may have been hindered by the time required for bacteria to be released and prime plant defences, i.e. aphids will have initiated feeding before defences are primed (Kim & Felton 2013). However, we observed significant suppression of aphids by *B. subtilis* when plants were inoculated after initial aphid infestation. This result is driven by a stronger reduction in offspring production in the latter half of the experiment. This contrasts with the results when inoculation occurred with aphid infestation, where we observed a non-significant increase in aphid reproduction in the latter part of the experiment. We cannot explain these differences from our current data, but we suspect this is related to fine-tuning of plant defence responses to aphid attack (Thompson & Goggin 2006) as well as establishing the rhizobacteria symbiosis (relevant to defence and nutritional pathways) (van de Mortel *et al*. 2012). Work in other systems indicates potential timing or context-dependency of rhizobacteria effects resulting in induced-susceptibility (Pineda *et al*. 2012) or induced-resistance (Pandharikar *et al*. 2020) by the same strain of *Pseudomonas simiae*. Future work focused on understanding plant responses at the mechanistic level will provide information to support our hypotheses further.

Our results confirmed previous work (Trivedi & Pandey 2008) on the longer-term storage and recovery of bacteria from alginate beads. We included additional nutrient broth in our formulation, and were able to recover comparable (if not higher) amounts of viable bacteria after eight months of storage at 4 °C. The diversity of uses for alginate bead inoculations is valuable for use in empirical experiments, particularly when used as a standalone inoculation (Hernández-Montiel *et al*. 2017) as opposed to seed coatings (Berninger, Mitter & Preininger 2016). Several benefits of larger beads include targeting the timing, placement, and concentration of inoculation. We can use multiple beads each containing different bacteria, or beads containing multiple strains, to study bacteria community interactions also incorporating spatial and temporal variables to disentangle composition and function of microbiomes (Song *et al*. 2020). We have further used these beads for large-scale inoculation of field trials (Zytynska, unpublished). This also allows for the control of time, location, and composition of inoculations within field trials. The use of water-dominated inoculant could also be beneficial, as it drenches the soil at time of addition to enable quicker breakdown of beads; however, we would also suggest to apply these just before a rainfall for optimal results. Alternative filler products used for seed coatings can contain compounds such as lime, talc, silica sand, gum arabic and various other, which could skew experimental conditions (Afzal *et al*. 2020) or alter microbiome responses (Ajeng *et al*. 2020; Brtnicky *et al*. 2021).

In conclusion, we show that the use of alginate bead encapsulation of plant beneficial rhizobacteria provides more consistent responses in the plants than seed soaking or dual-sowing seeds and beads. The greatest suppression of aphids on barley plants occurred when beads were added to the plant roots at root emergence and before the plants were infested by aphid pests. The benefits of using alginate beads in empirical research includes precision timing and location of inoculation, as well as the ability to study bacteria-bacterial interaction with temporal and spatial variables. Field-trial inoculations using alginate beads benefit from the long-term viability (shelf-life) of the encapsulated bacteria. While there is still substantial research and development needed for wider application and use of bacterial beads, it is essential that we continue to optimise the timing and application methods.

## Acknowledgements

The data presented was generated as part of a MRes (Blenkinsopp) and an undergraduate project (Henderson) at The University of Liverpool, UK and was supported by a BBSRC David Phillips Fellowship BB/S010556/1 (Zytynska).

## Author contributions

All authors contributed to experimental design, set up and data collection. Data were first analysed by SB (bead formulation and Expt 1) and MH (Expt 2), refined by SEZ. The first manuscript draft was written by SB, with input from MH, with the final version edited by SEZ.

